# Comparative assessment of machine learning algorithms to predict severity of disease in COVID-19 patients based on eight cofactors

**DOI:** 10.1101/2024.10.11.617884

**Authors:** Tanvi S. Patel, Daxesh P. Patel, Hoda El-Sayed, Mallika Sanyal, Pranav S. Shrivastav

**Author notes:** **Correspondence author** Pranav S. Shrivastav; (M).

## Abstract

Machine learning is one of the important tools to diagnose and predict the diseased state accurately and effectively. The COVID-19 pandemic caused due to severe acute respiratory syndrome coronavirus 2 (SARS-CoV-2) has become one of the most researched healthcare topics worldwide. Machine learning algorithms can find efficient and reliable ways to predict the COVID-19 from vast amounts of existing health care data, allowing faster, effective, and more accurate diagnosis with lower risk based on the symptoms. Based on the countrywide data published by the Israeli Ministry of Health, we propose a system that detects COVID-19 instances using simple variables. The COVID-19 dataset used in the study consisted of 278848 patients samples with five different symptoms, namely cough, fever, sore throat, shortness of breath, and headache, apart from other basic information like age, gender, and test indication excluding confirmed COVID-19 result. The data was analyzed using traditional supervised machine learning algorithms namely, Decision tree, Support vector machine, Random Forest, Logistic regression, k-nearest neighbor, and Naive Bayes based on eight cofactors with high accuracy rate (≥ 0.9450). Apart from Support vector machine, all other algorithms displayed better performance based on the AUC score calculated using the receiver operator characteristic (ROC) curve. This study also highlights the significant differences between precision, recall and accuracy for each model.

Graphical Abstract

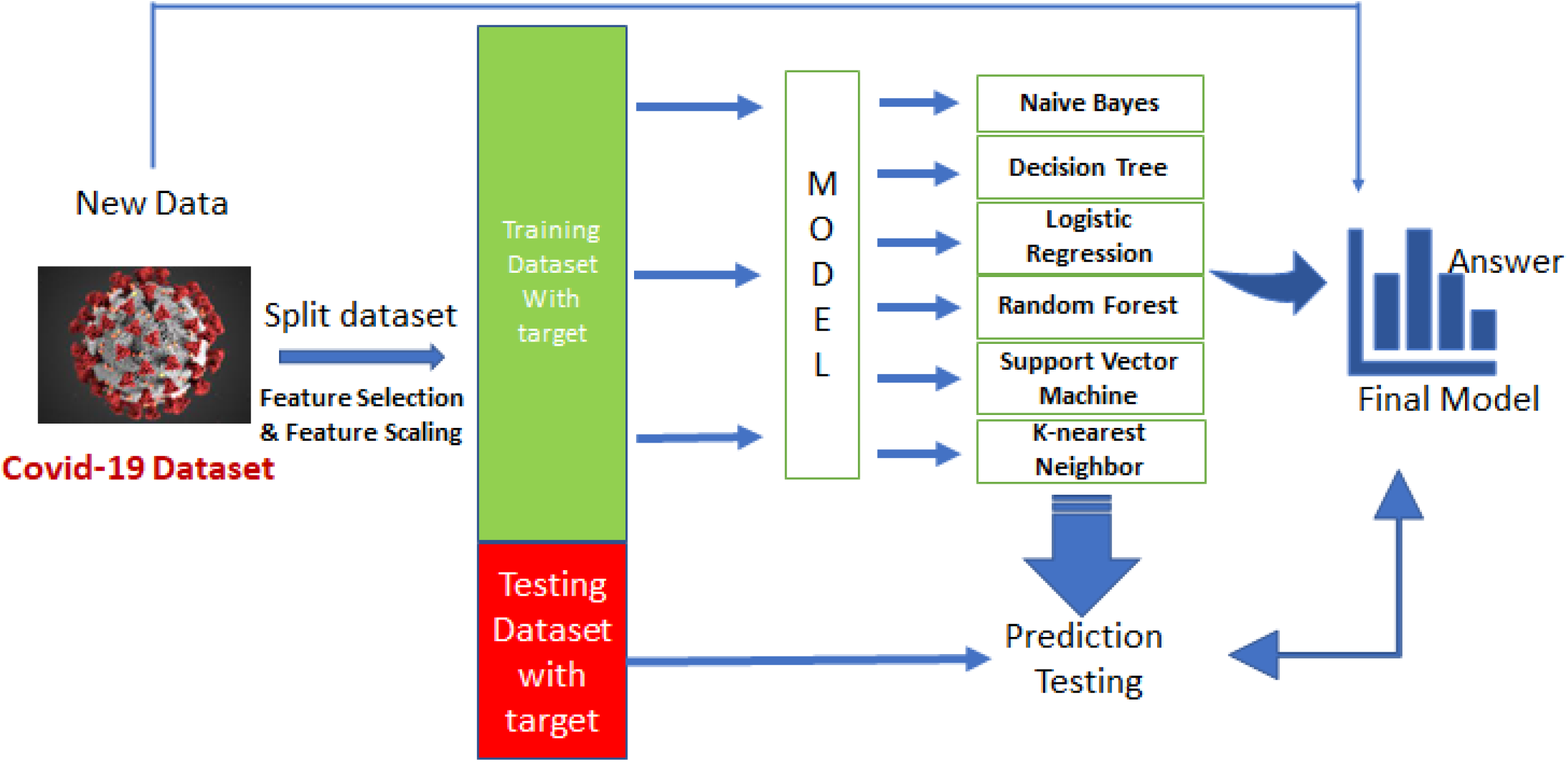

## 1. Introduction

There has been a global concern to control and minimize the effect of severe acute respiratory syndrome coronavirus-2 (SARS-CoV-2) which has severely impacted healthcare services worldwide. On December 31, 2019, the World Health Organization (WHO) received the first report of coronavirus outbreak [1–3] and recognized this virus as novel coronavirus "2019-nCoV" (COVID-19) on January 12, 2020. The International Committee on Taxonomy of Viruses (ICTV) later described it as SARS-CoV-2, while WHO labeled this viral epidemic a public health emergency of worldwide concern on January 30, 2020. Since SARS-CoV-2 has spread swiftly in more than 220 countries, WHO proclaimed COVID-19 a global pandemic [4, 5]. More than 490 million persons have been infected with coronavirus as on March 27, 2022. Different countries have enforced several measures to prevent the spread of this deadly infection by controlling social interactions, and quarantine of patients under strict guidelines. Although the fatality rate is low, it is responsible for about 5.7 million deaths worldwide. The recovery rate has increased, and the mortality rate has decreased since the COVID-19 immunization began in January 2021 [6–8]. However, due to some unique symptoms that vary from patient to patients it has become difficult to give proper treatment after diagnosis. Besides, sizeable number of positive COVID-19 cases have been observed even after immunization. The Centers for Disease Control and Prevention (CDC) and other public health institutions worldwide keep a track of all COVID-19 viral variations [9, 10]. Although the Research of University of California proved that the most prevalent symptoms of COVID-19 were cough and fever, shortness of breath develops in some patients, chills, trembling (shaking), runny nose, sore throat, muscle discomfort, headache, lethargy, and loss of smell or taste are also observed [11, 12].

As COVID-19 is associated with different clinical manifestations, forecasting mortality, disease severity and outcome can provide crucial information for critically ill patients. Machine learning (ML) algorithms can help in developing prediction models and thereby reduce the complexity of clinical traits. They have been successfully used as diagnostic tool or predictive model in health care and medical informatics [13]. The major goal of these models is to assist the medical industry in predicting patients with symptoms and initiating early therapy, as well as to provide fast and accurate results. Several researchers have studied different ML algorithms for early disease prediction, criticality and chances of survivability using existing clinical data. Aljameel et al. [14] developed prediction model for the early identification of COVID-19 patient’s outcome based on patients’ characteristics. They performed the study using 287 COVID-19 patients samples using logistic regression (LR), random forest (RF), and extreme gradient boosting (XGB). Similar studies have been carried out using XGB model to predict criticality and chances of survival for patients with severe COVID-19 infection [15, 16] and mortality risk [17]. Support vector machine (SVM) algorithm has also been applied to predict severity of the symptoms of COVID-19 patients [18, 19]. An et al. [20] developed a model to predict the mortality of COVID-19 patients using different ML algorithms such as Least absolute shrinkage and selection operator (LASSO), SVM, RF, and k-nearest neighbor (KNN).

In the present work we have tried to predict the severity of COVID 19 in patients based on the symptoms such as cough, fever, sore throat, shortness of breath and headache for building ML prediction model. The study was performed using publicly available 278848 sample data from Ministry of Health, Government of Israel. Following preprocessing, we obtained 136294 patient samples with no missing data and after handling incomplete information. Out of which 70 % samples were used for training and 30 % for testing dataset, respectively. Zoabi and co-workers [21] proposed a ML model that predicted a positive SARS-CoV-2 infection (via RT-PCR assay of nasopharyngeal swab) by asking eight basic questions. Their model was trained on records from 51,831 tested individuals, while the test set contained 47,401 tested individuals. In our work, eight features in the dataset were studied using six ML algorithms, namely Decision tree (DT), SVM, RF, LR, KNN and Naïve Bayes (NB). The sensitivity/recall, precision, accuracy, F1 score and area under the curve (AUC) of the algorithms was compared to find any significant difference (P < 0.05) between the models.

## 2. Dataset source and machine learning algorithms

### 2.1 Dataset used

The Ministry of Health, Government of Israel made available the data of all people who were tested via RT-PCR assay using nasopharyngeal samples. During the early months of the COVID-19 pandemic in Israel, all COVID-19 diagnostic laboratory tests were performed according to Ministry of Health guidelines [22–24]. The report on ML-based prediction of COVID-19 utilized this data by applying gradient-boosting ML model with DT [22]. To develop the models in our proposed approach, we selected three ML techniques and compared their results. **Table 1** shows the description of raw dataset.

**Table 1.** COVID-19 prediction dataset.

| Sr. No. | Attribute/ Feature | Not-Null Count | Data type | Description |
| --- | --- | --- | --- | --- |
| 1 | Test date | 278848 | Object | Between 11<sup>th</sup> March 2020 to 30<sup>th</sup> April 2020 |
| 2 | Cough | 278848 | Object | 0-No, 1- Yes, None - patients who said nothing |
| 3 | Fever | 278848 | Object | 0-No, 1- Yes, None - patients who said nothing |
| 4 | Sore throat | 278848 | Object | 0-No, 1- Yes, None - patients who said nothing |
| 5 | Shortness of breath | 278848 | Object | 0-No, 1- Yes, None - patients who said nothing |
| 6 | Headache | 278848 | Object | 0-No, 1- Yes, None - patients who said nothing |
| 7 | Corona result | 278848 | Object | 0-Negative, 1- Positive, None - patients who did not prefer to answer |
| 8 | Age 60 and above | 278848 | Object | 0-No, 1- Yes, None - patients who did not prefer to answer |
| 9 | Gender | 278848 | Object | Female, Male and None |
| 10 | Test indication | 278848 | Object | Others, confirmed contact with COVID-19 patient, coming from abroad |

### 2.2 Machine learning models

In supervised ML new information can be obtained from the previously learned data to predict future events. Here, the algorithm is allowed to learn from a data set with pre-defined labels (supervisor) and the output specified by the learning system can be compared with the real output [25, 26]. Besides, it can also provide feedback on the accuracy of the prediction during the training process. Based on the Ministry of Health, Government of Israel dataset, we applied supervised learning algorithms such as DT, SVM, RF, LR, KNN and NB to compare their performance and forecast severity of COVID-19 patients in this study. DT generates classification or regression models in the form of a tree structure. Further, RF and SVM works for both classification and regression problems, but this study showed that DT outperforms SVM in categorization classified dataset prediction. In DT, a dataset is incrementally broken down into smaller portions while concurrently creating a decision tree. The data is divided through the hyperplane in SVM, and the results are obtained. **Figure 1** shows the workflow and the algorithms used in the models.

**Figure 1.**
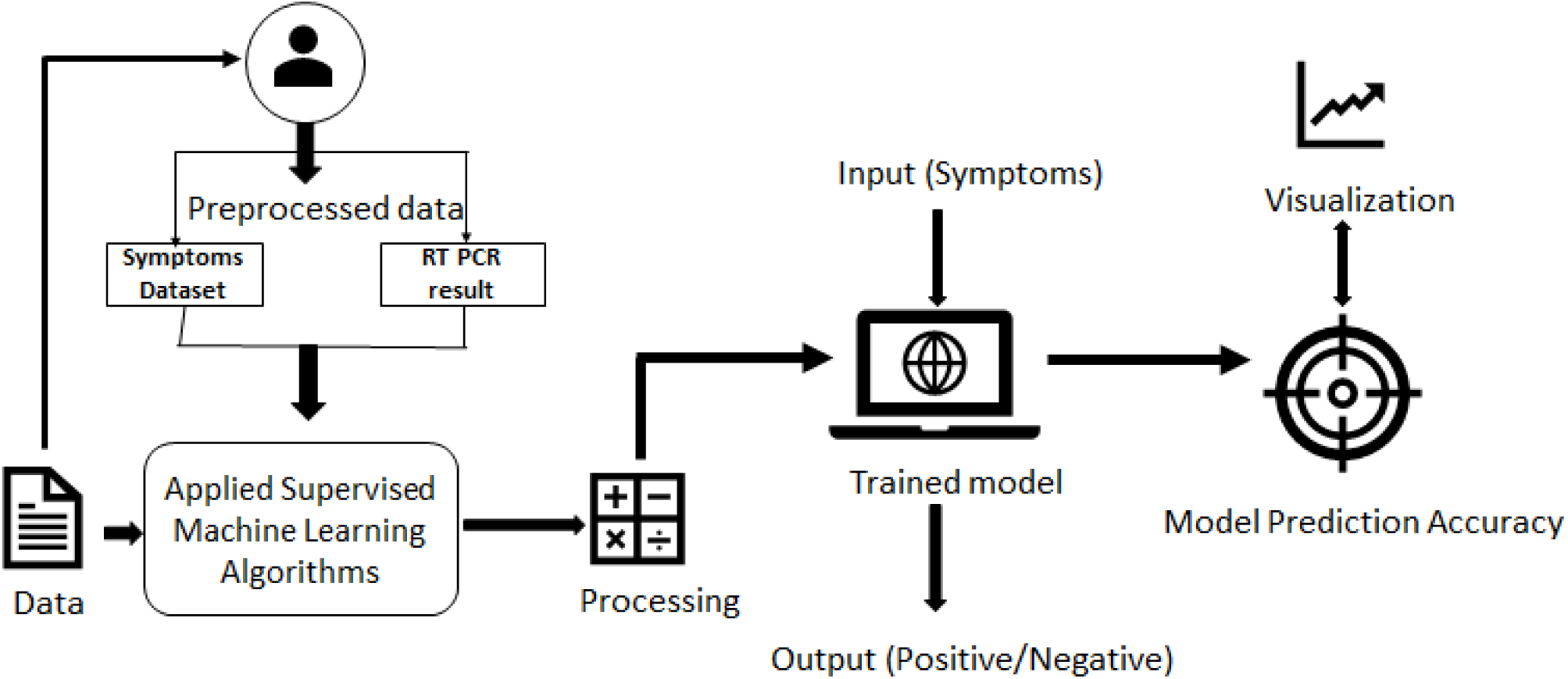
COVID-19 detection using supervised machine learning algorithms.

### 2.3 Framework and data preprocessing

In the development of prediction model, python programming was applied to create ML models. Python is a general-purpose and the most popular programming language among the data scientists and can do a variety of complicated ML tasks. All the python libraries used in this investigation worked better with python, including pandas, NumPy, Matplotlib, and Scikit-learn. First, we changed our csv data file to pandas, which is faster and has a much smarter ML backbone than .csv. Pandas can also be used to analyze and visualize data to spot trends and patterns. Further, using Python core programming, the data was preprocessed to change the data types from object to numerical and categorical. Here it is worth mentioning that the missing data was carefully treated. The dataset was divided into 70 % training and 30 % test. The dataset was encoded and transformed into categorical data using scikit-learn to apply ML algorithms.

To create the prediction model, we used five COVID-19 symptoms from the dataset and matplotlib was utilized for all data visualization. In the training dataset, we used three ML algorithms and predicted the results using the test dataset and utilized target prediction with test prediction to determine the model’s accuracy after fitting datasets into various algorithms [27]. The recall rate, precision, and AUC of all the models were assessed after determining the model accuracy. Each model has a separate and unique recall, precision, and AUC score, therefore we compared them to see which one works best and if there is any significant difference (P < 0.05) in accuracy and AUC score using the receiver operator characteristic (ROC) curve. There were 278848 total patients’ data in the raw dataset and after preprocessing, the training and testing steps were applied to the remaining 136294 dataset. After splitting the dataset there were 95405 patient records that were utilized to train the dataset and fit various models. Following the fitting of the model, 40889 patients testing dataset was utilized to evaluate the model and determine the various scores for the models. Considering all the features followed a Gaussian distribution, Gaussian NB was used in the study. In RF classifier, we used ‘n’ estimators’ values of 10. DT classifier for DT and SV classifier and a linear kernel was utilized for SVM. For KNN, the number of ‘n’ neighbors (185) was calculated by taking the square root of 136294 and dividing by 2.

### 2.4 Analysis and performance evaluation

The COVID-19 dataset comprised of ten attributes, including the basic information, test date and COVID-19 result (**Table 1**) [28]. Out of 278848 total patients’ samples in the dataset, there were 130158 females, 129127 males, and for 19563 patients there was no mention of the gender. From the patients who were tested for RT-PCR assay, 260227 patients were found to be corona virus negative, 14729 patients were corona virus positive, while 3892 patients revealed neither negative nor positive results. The data given is from 11^th^ March 2020 to 30^th^ April 2020, which was the initial period of the COVID-19. **Table S1** shows the samples used during each step of the data preprocessing. After preprocessing the data, we had 136,294 samples, which were divided into 70 percent training and 30 percent testing, yielding 95405 samples in the train dataset, while 40889 samples were used to test the models.

Confusion matrix was obtained after applying the six ML algorithms to the training dataset, to find the accuracy of different models as well as to determine the model accuracy, recall, and precision rate using basic math calculations. A 2 x 2 matrix was used to evaluate the performance of a classification model in case of positive and negative target classes. The matrix compares the actual goal values to the ML model’s prediction [29]. We utilized scikit-learn to extract the confusion matrix in the model, and then applied the confusion matrix to that mode to determine the accuracy, sensitivity, F1 weighted average score of the true positive, and ratio of correctly detected positive class. The rows of the confusion matrix indicate instances of the real class, whereas the columns represent expected classes. The true positive (TP) and true negative (TN) values were defined as correctly predicted positive and negative samples, respectively. While false negative (FN) and false positive (FP) were incorrectly predicted positive and negative samples, respectively. The method sensitivity/recall, precision, accuracy and F1 score (weighted value of the precision and sensitivity) were expressed using the following equations

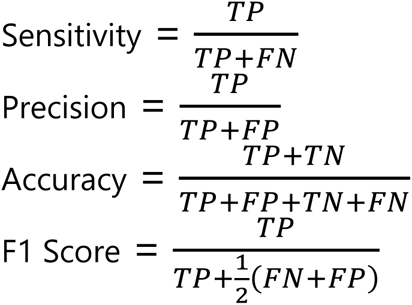

## 3. Results and discussion

The bar chart (**Figure S1**) illustrates the importance of different features studied in the present work. The values obtained after calculating the dependency of features with the target values for cough, fever, sore throat, shortness of breath, headache, age 60 and above, gender, and test indication were 0.0208, 0.0341, 0.0234, 0.0159, 0.0360, 0.0004, 0.0077, and 0.1018, respectively. The test indication showed a significant impact on the target variable.

Violin plot which gives a visual representation of entire distribution of the numerical data was used in the present work. A violin plot is a fusion of a box plot and a kernel density plot, which shows the data’s peak and depth at various points. Contrary to a box plot that can only show summary statistics, violin plots depict summary statistics and the density of each variable. It was used to depict the distribution of numerical data categorized as positive or negative in COVID-19. **Figure 2** shows the violin plot for the eight cofactors that affect COVID- 19 positive and negative patients. Findings for shortness of breath on COVID-19 shows that most patients were negative, as the data distribution is greater at the corona negative result (compared to the positive result) around zero and the mean and median value are also at the same point (**Figure 2A**). However, the data distribution for shortness of breath, headache and sore throat symptoms is minimal at corona positive case at point 1 (**Figure 2A-C**). The data distribution at negative and positive result is similar for the violin plots of shortness of breath and sore throat. In contrast to shortness of breath, headache, and sore throat, the positive corona result at point 1 is quite noticeable for cough and fever symptoms (**Figure 2D and E**). In the gender violin plot, the positive male gender has a little greater depth than the positive female gender (**Figure 2F**). The plot of test indications reveals that contact with confirmed patients is mainly positive (**Figure 2G**). For people above the age of 60, the violin plot indicates that the ratio of tested positive and tested negative is similar (**Figure 2H**).

**Figure 2.**
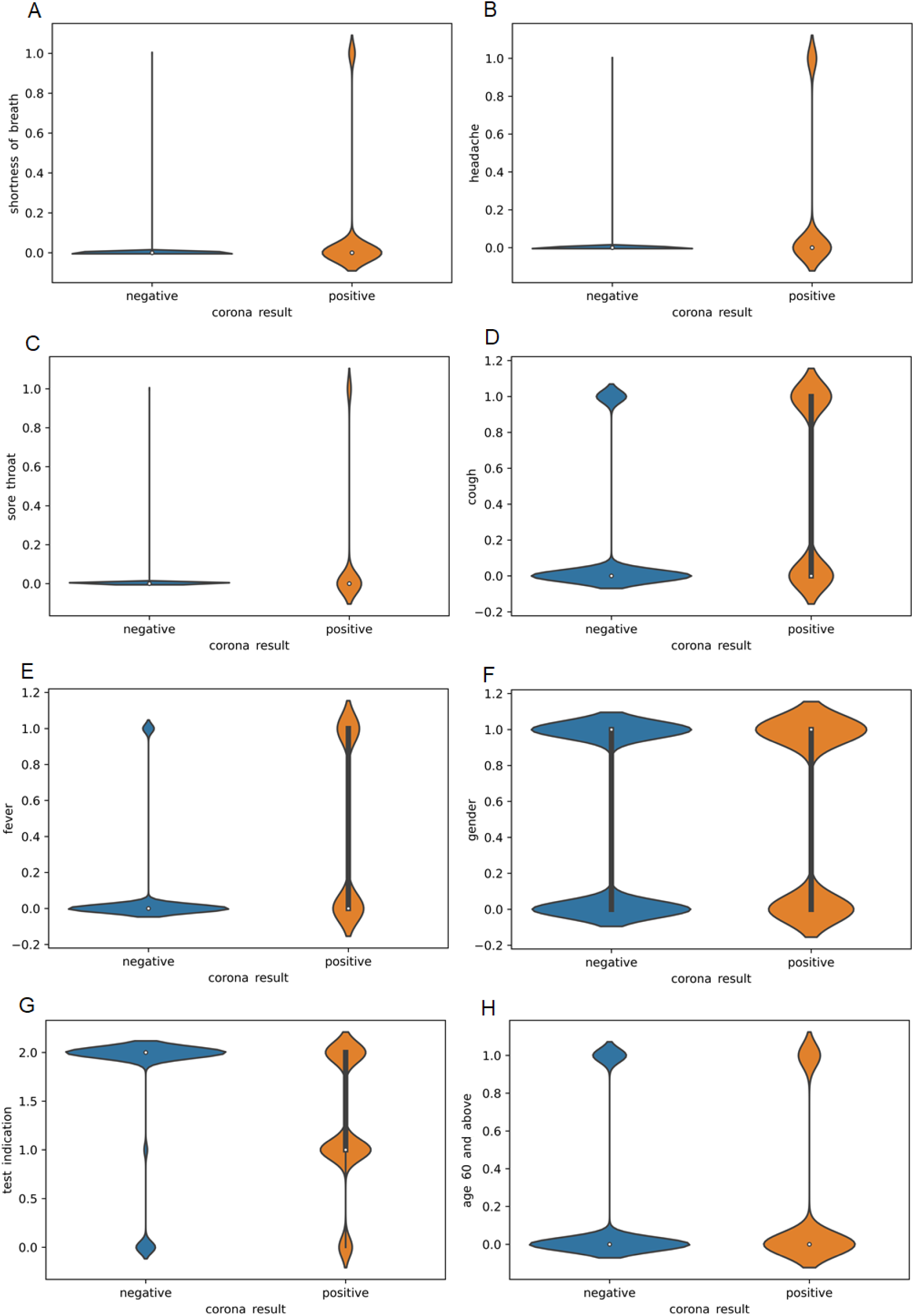
Violin plots for eight cofactors that affect COVID-19 patients results, (A) shortness of breath, (B) headache, (C) sore throat, (D) cough, (E) fever, (F) gender (1 = male, 0 = female), (G) test indication (2-others, 1-confirmed contact with COVID-19 patient, 0-coming from abroad) and (H) age 60 and above.

Confusion matrix was used to calculate sensitivity, precision, accuracy and F1 score. **Figure 3** depicts the confusion matrix for the various models showing the true positive values in the top left corner, false positive values in the top right corner, false negative values in the bottom left corner and true negative values in the bottom right corner [30, 31]. For LR and SVM the confusion matrix was identical, the TP, FP, FN, and TN values were 37651, 65, 2183 and 990, respectively. Such similarity was also found for DT and RF, and the corresponding values were 37322, 394, 1352 and 1815, respectively. For NB, the confusion matrix values were 37680, 66, 2172 and 971, while for KNN the values were 37390, 326, 1709 and 1464, respectively. The random state was kept at one for all the models. As a result, the sensitivity, precision, accuracy and F1 score were the same for LR & SVM, and DT & RF algorithms. **Table 2** displays the outcomes of all models, including recall, precision, accuracy, F1 score, and AUC. We also evaluated the significant difference (P < 0.05) between all the models by comparing models’ AUC, sensitivity, precision, accuracy, and F1 score (**Table 3**). There were no significant differences in accuracy between all the models. While comparing the model efficiency using the AUC values obtained from the ROC curves (**Figure 4**), the results demonstrate significant difference between SVM and other algorithms, with no significant change among the other models. For various thresholds, the precision-recall curve depicts the tradeoff between precision and recall (**Figure 5**). The AUC value in RF model was 0.91, while the average precision in the precision recall curve was 0.71. Thus, it can be deduced that the RF model is associated with low false positive and false negative rates. Besides, a high AUC indicates a high recall as well as a high precision. The average precision rate for DT and KNN was 0.70 and 0.65, respectively, and have low positive and false negative as compared to LR, SVM, and NB, which have precision values of 0.52, 0.45, and 0.51, respectively.

**Figure 3.**
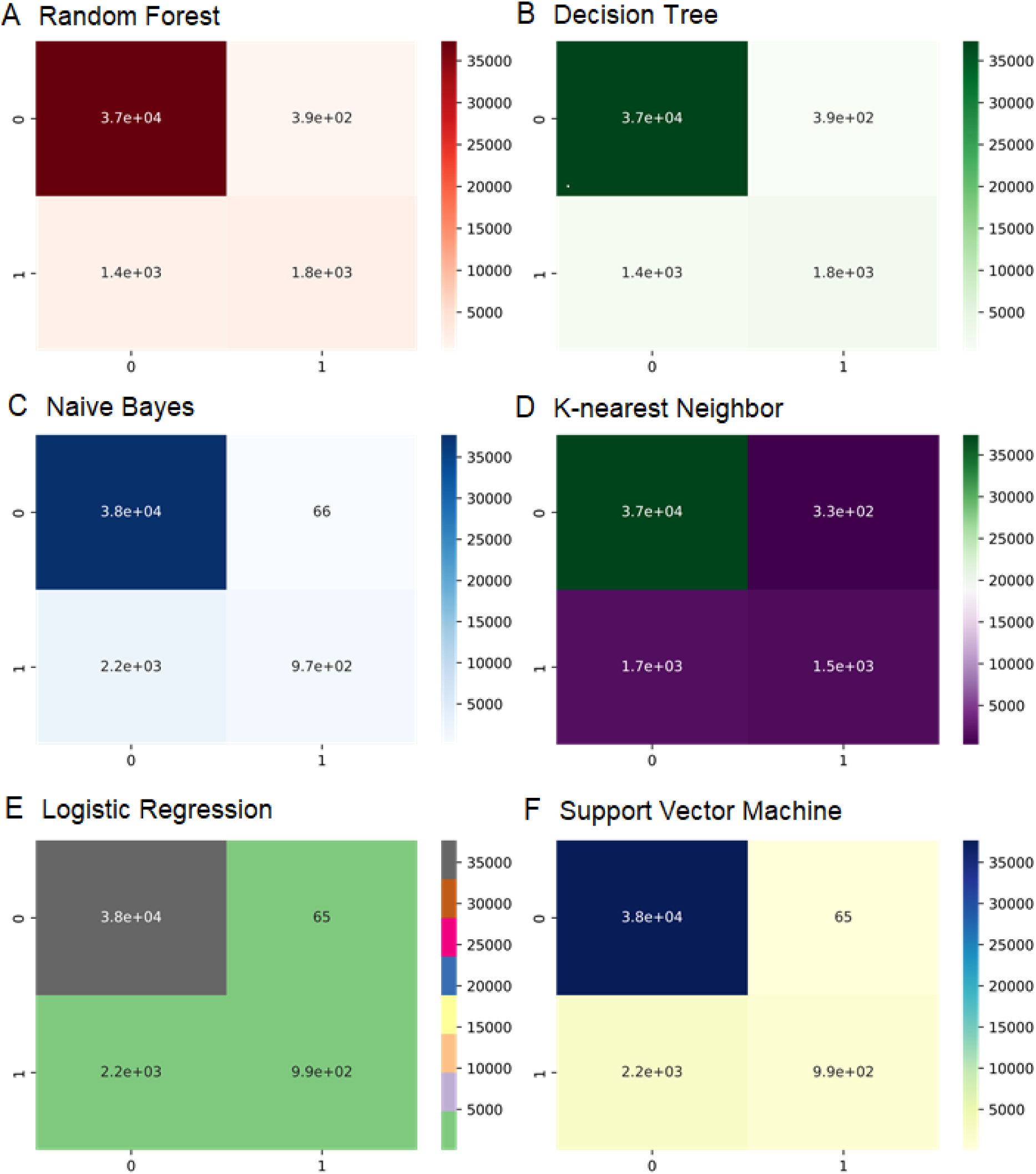
Experiment of confusion matrices for (A) Random Forest, (B) Decision tree, (C) Naïve Bayes, (D) k-nearest neighbor, (E) Logistic Regression and (F) Support vector machine algorithms to calculate the results.

**Figure 4.**
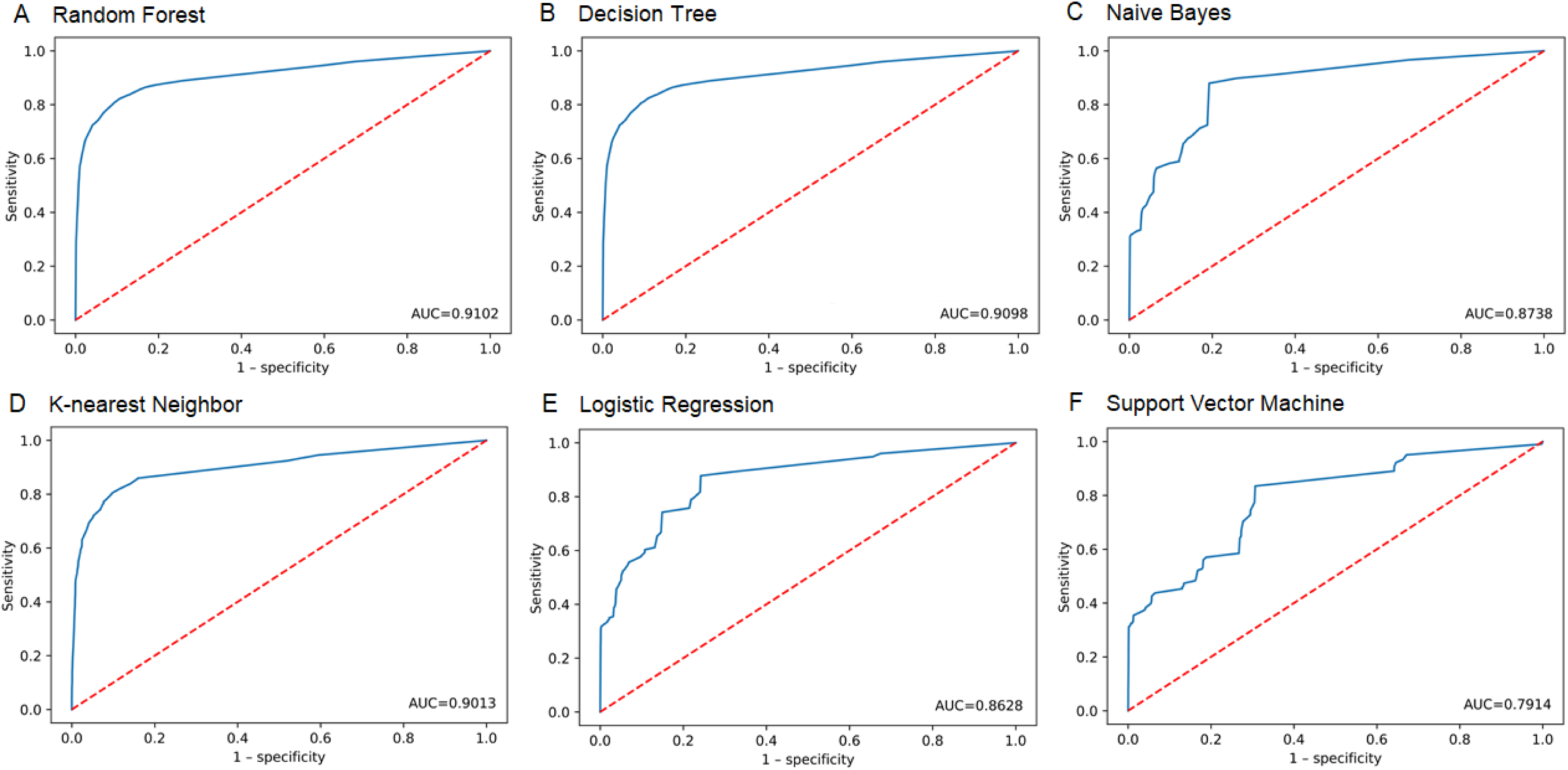
Model performance using receiver operating characteristics (ROC) curves for (A) Random Forest, (B) Decision tree, (C) Naïve Bayes, (D) k-nearest neighbor, (E) Logistic Regression and (F) Support vector machine algorithms.

**Figure 5.**
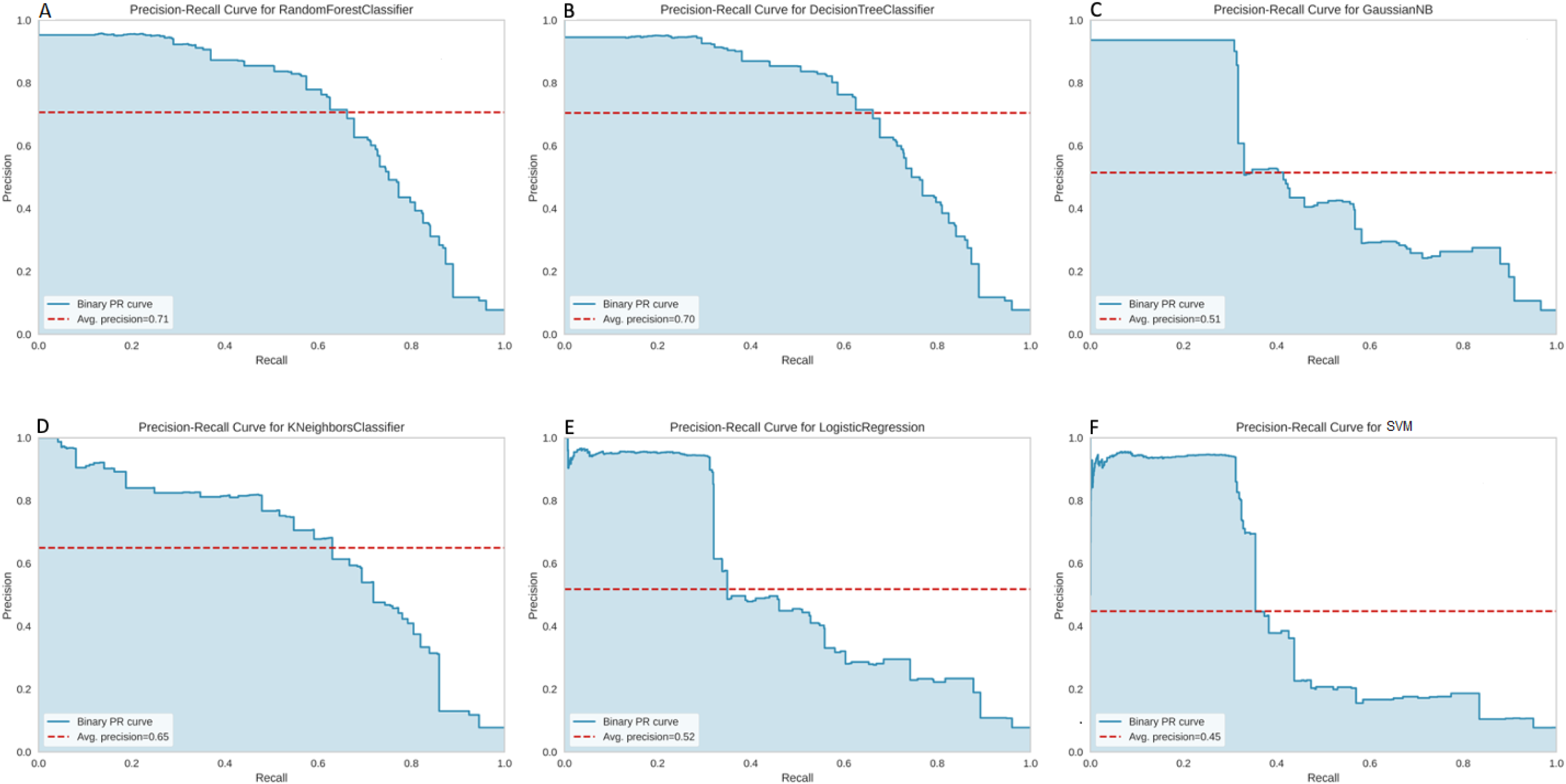
Model performance using precision recall curves for (A) Random Forest, (B) Decision tree, (C) Naïve Bayes, (D) k-nearest neighbor, (E) Logistic Regression and (F) Support vector machine algorithms.

**Table 2.** Performance of the supervised machine learning models.

| Model | Accuracy | Error | Precision | Sensitivity/<br>Recall | F1 score | AUC |
| --- | --- | --- | --- | --- | --- | --- |
| Naive Bayes | 0.9453 | 0.0547 | 0.9364 | 0.3089 | 0.9322 | 0.8738 |
| Support vector machine | 0.9450 | 0.0550 | 0.9384 | 0.3120 | 0.9320 | 0.7914 |
| Decision Tree | 0.9572 | 0.0428 | 0.8216 | 0.5720 | 0.9536 | 0.9099 |
| Random Forest | 0.9572 | 0.0428 | 0.8216 | 0.5720 | 0.9536 | 0.9102 |
| k-Nearest neighbor | 0.9502 | 0.0498 | 0.8179 | 0.4614 | 0.9437 | 0.9013 |
| Logistic Regression | 0.9450 | 0.0550 | 0.9384 | 0.3120 | 0.9320 | 0.8628 |

**Table 3.**
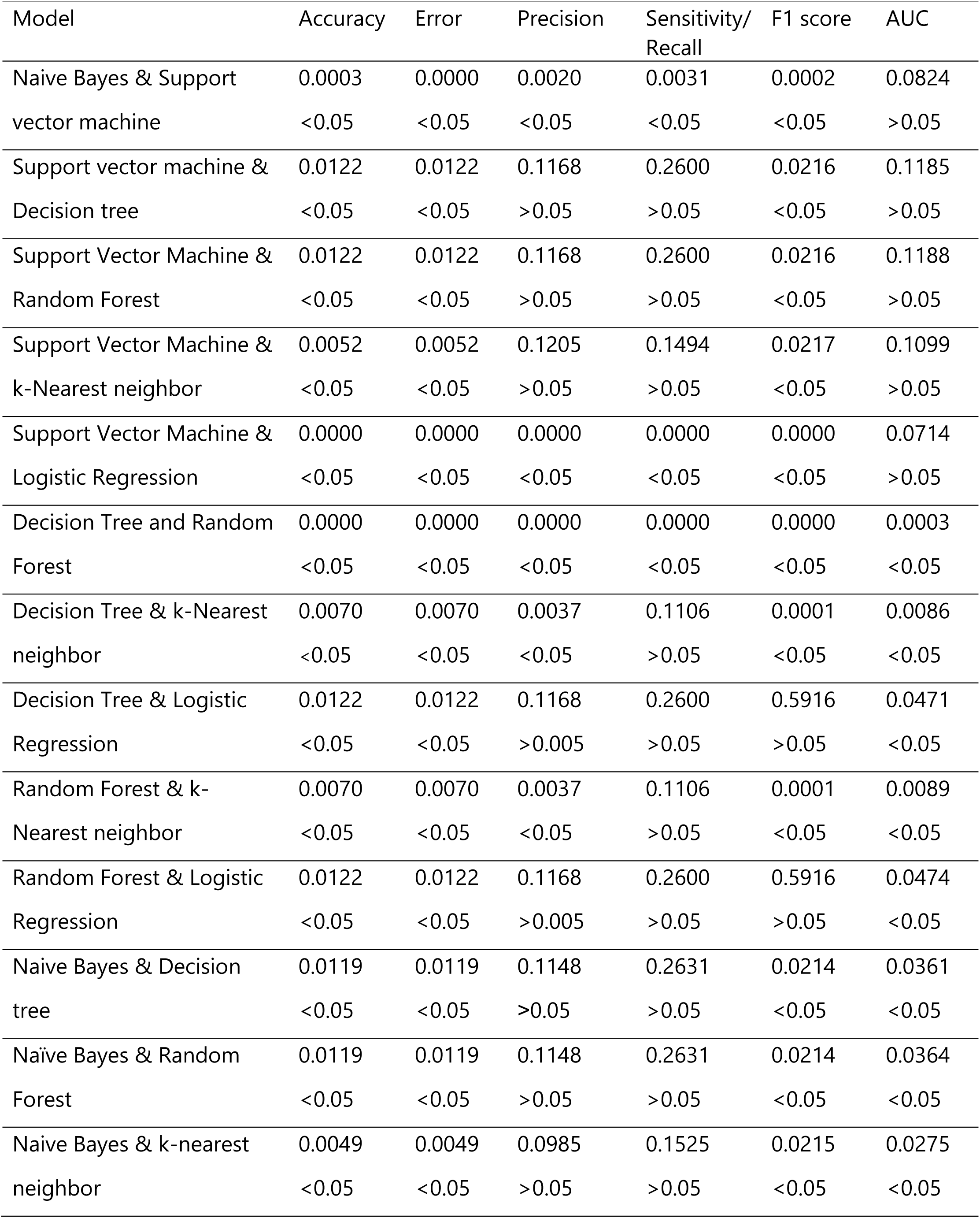
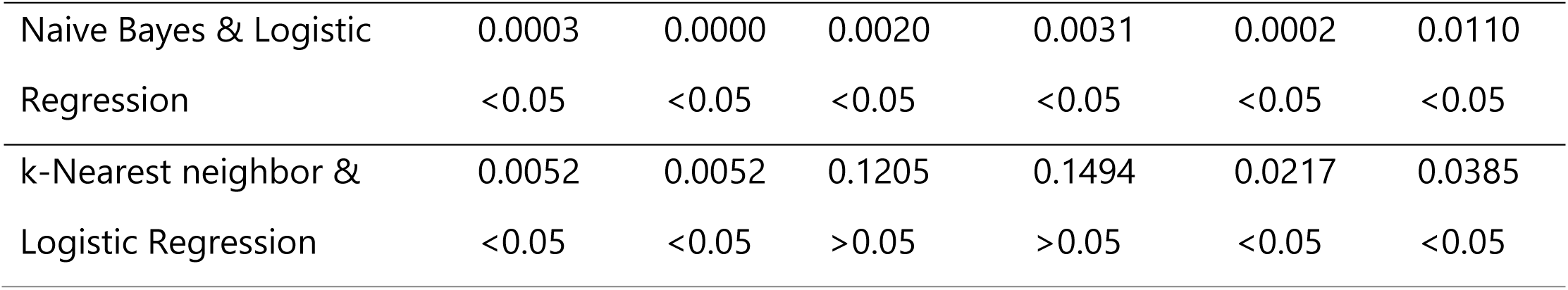
Comparison of models with significance level.

Although there are several similar works reported in literature using different prediction models with accuracy ranging from 0.71-0.95, the dataset used have been limited (115-8000) [14–20]. Only one report established an RF ML approach that employed 51,831 samples for training and 47,401 as test set using Israeli Ministry of Health publicly released data (Zoabi et al., 2021). The AUC values obtained (0.91) were comparable (except SVM) but the accuracy of the present models was also higher (˃ 0.95) compared to their work. They used biased and unbiased to show the importance of features and how they affect the model in that work. However, in the proposed model, we used all the features on the same datasets and obtained higher AUC and accuracy than the previous model on the same algorithm (RF) used in their work.

## 4. Conclusion

In this study we utilized six ML algorithms to detect severity of COVID-19 based on symptoms. All the supervised ML algorithms applied on 278848 patient records demonstrated promising accuracy of ≥ 0.9450. The data suggested good correlations with the COVID-19 severeness. Comparing the difference between the AUC values calculated using the ROC showed no substantial difference, except between SVM and all other algorithms. Despite these advantages, the models can have inherent limitations as we relied on the data reported by the Israeli Ministry of Health, which had some missing information regarding some of the attributes studied. Nevertheless, we believe the proposed models can be useful especially for resource limited setups and may help health care personnel to take quick and reliable decisions for treatment of COVID-19 patients.

## Author’s contributions

**TSP** devised and designed the study, wrote the study protocol, contributed to data collection and analyses.

**DPP** contributed to the study idea and protocol and did analyses.

**HES** contributed to the study idea and protocol.

**MS** contributed substantially towards writing the draft manuscript.

**PSS** contributed towards critically revising, editing the manuscript and supervision. Further, all authors read and approved the final manuscript for submission

## Availability of data and materials

There is no data or material to report.

## Acknowledgments

We would like to thank Department of Computer Science, Bowie State University and Department of Chemistry, Gujarat University for supporting this work.

